# Longitudinal characterization of humoral and cellular immunity in hospitalized COVID-19 patients reveal immune persistence up to 9 months after infection

**DOI:** 10.1101/2021.03.17.435581

**Authors:** John Tyler Sandberg, Renata Varnaitė, Wanda Christ, Puran Chen, Jagadeeswara R. Muvva, Kimia T. Maleki, Marina García, Majda Dzidic, Elin Folkesson, Magdalena Skagerberg, Gustaf Ahlén, Lars Frelin, Matti Sällberg, The Karolinska COVID-19 Study Group, Lars I. Eriksson, Olav Rooyackers, Anders Sönnerborg, Marcus Buggert, Niklas K. Björkström, Soo Aleman, Kristoffer Strålin, Jonas Klingström, Hans-Gustaf Ljunggren, Kim Blom, Sara Gredmark-Russ

## Abstract

**Background:** Insights into early, specific humoral and cellular responses to infection with SARS-CoV-2, as well as the persistence and magnitude of resulting immune memory is important amidst the ongoing pandemic. The combination of humoral and cellular immunity will most likely contribute to protection from reinfection or severe disease.

**Methods:** Here, we conducted a longitudinal study on hospitalized moderate and severe COVID-19 patients from the acute phase of disease into convalescence at five- and nine-months post symptom onset. Utilizing flow cytometry, serological assays as well as B cell and T cell FluoroSpot assays, we assessed the magnitude and specificity of humoral and cellular immune memory during and after human SARS-CoV-2 infection.

**Findings:** During acute COVID-19, we observed an increase in germinal center activity, a substantial expansion of antibody-secreting cells, and the generation of SARS-CoV-2-neutralizing antibodies. Despite gradually decreasing antibody levels, we show persistent, neutralizing antibody titers as well as robust specific memory B cell responses and polyfunctional T cell responses at five- and nine-months after symptom onset in both moderate and severe COVID-19 patients. Long-term SARS-CoV-2 specific responses were marked by preferential targeting of spike over nucleocapsid protein.

**Conclusions:** Our findings describe the initiation and, importantly, persistence of cellular and humoral SARS-CoV-2 specific immunological memory in hospitalized COVID-19 patients long after recovery, likely contributing towards protection against reinfection.

## Introduction

Severe acute respiratory syndrome coronavirus 2 (SARS-CoV-2), the causative agent of coronavirus disease 2019 (COVID-19), emerged in late 2019 and has since led to a pandemic resulting in deaths of more than 2 million people within just one year^1,2^. COVID-19 is primarily a respiratory disease with severity ranging from asymptomatic or mild infection to severe symptoms requiring hospitalization with oxygen supplementation or mechanical ventilation. The development of adaptive immune responses, encompassing neutralizing antibodies, B cells and T cells, is crucial to control and clear viral infections^3^. Studying early cellular and humoral responses to infection with SARS-CoV-2 can give insights into the development of immune memory. These early responses can be assessed by germinal center activity via plasma CXCL13 levels and activated circulating T follicular helper cells (cTfh) frequencies as surrogate markers^4–7^, as well as antigen specific antibody-secreting cell (ASC) expansion^8–11^. Once infection is cleared, neutralizing antibodies and antigenspecific memory B cells and T cells play an important role in preventing reinfection. Recent reports suggest that cellular and humoral immunity to SARS-CoV-2 lasts up to at least 8 months post-infection^12,13^. However, studies of related viruses including SARS-CoV-1 and Middle Eastern Respiratory Syndrome (MERS) coronavirus have shown that specific cellular memory persists longer than antibodies after infection^14,15^ and similar patterns may be expected for SARS-CoV-2. It is therefore important to characterize, in detail, the specificity and magnitude of adaptive immune responses throughout the course of disease and during recovery, in order to better understand the development and longevity of protective immunity.

Here, we provide an in-depth assessment of the initiation and persistence of antigen-specific humoral and cellular immune responses in hospitalized COVID-19 patients, suffering from moderate or severe disease. In this context, we report increased germinal center activity and ASC expansion during the acute phase followed by persistent SARS-CoV-2-specific humoral and cellular immunity. Neutralizing antibodies, memory B cells and polyfunctional memory T cells specific to SARS-CoV-2 were detectable in all COVID-19 patients in convalescence, regardless of disease severity, potentially providing long-term protection against reinfection.

## Results

### Clinical features and outcome of hospitalized COVID-19 patients

Twenty-six hospitalized COVID-19 patients were included in this study, ten of whom were treated at the infectious disease unit (IDU) and are referred to as moderate, while sixteen patients were treated at the intensive care unit (ICU) and are referred to as severe (Figure 1A; Table 1). The present patient cohorts were part of the Karolinska KI/K COVID-19 Immune Atlas project, where characterizations of other immune cell subsets during acute COVID-19 have been described^16–21^. When possible, these patients were sampled longitudinally at 5- and 9-months after symptom onset (Figure 1A; Table 1). Patients were predominantly male, with comparable age between moderate and severe groups (Figure 1B and C). Patients in both groups were also sampled at similar timepoints since symptom onset during the acute phase, with a median of 14 days for both groups (Figure 1D; Table 1). Patients treated in the ICU required longer supplemental oxygen treatment compared to moderate patients (a median of 2 days for moderate, and a median of 21 days for severe) (Figure 1E). Respiratory organ failure assessment score (SOFA-R) ranged from 1-2 for moderate and 2-4 for severe patients^22^. Patients showed considerable alterations from normal values in clinical chemistry parameters, including inflammatory and tissue damage markers, and significant differences were observed between the moderate and severe groups in several of those parameters during the acute phase of COVID-19 (Figure S1). Both patient groups presented with characteristic symptoms of COVID-19 and suffered from common co-morbidities (Figure 1F; Table 1). Four patients deceased, all of whom were in the severe group (Figure 1F; Table 1). The majority of patients followed up at 5-month convalescence had returned to normal levels in inflammatory and tissue damage markers (Figure S1).

**Figure 1.**
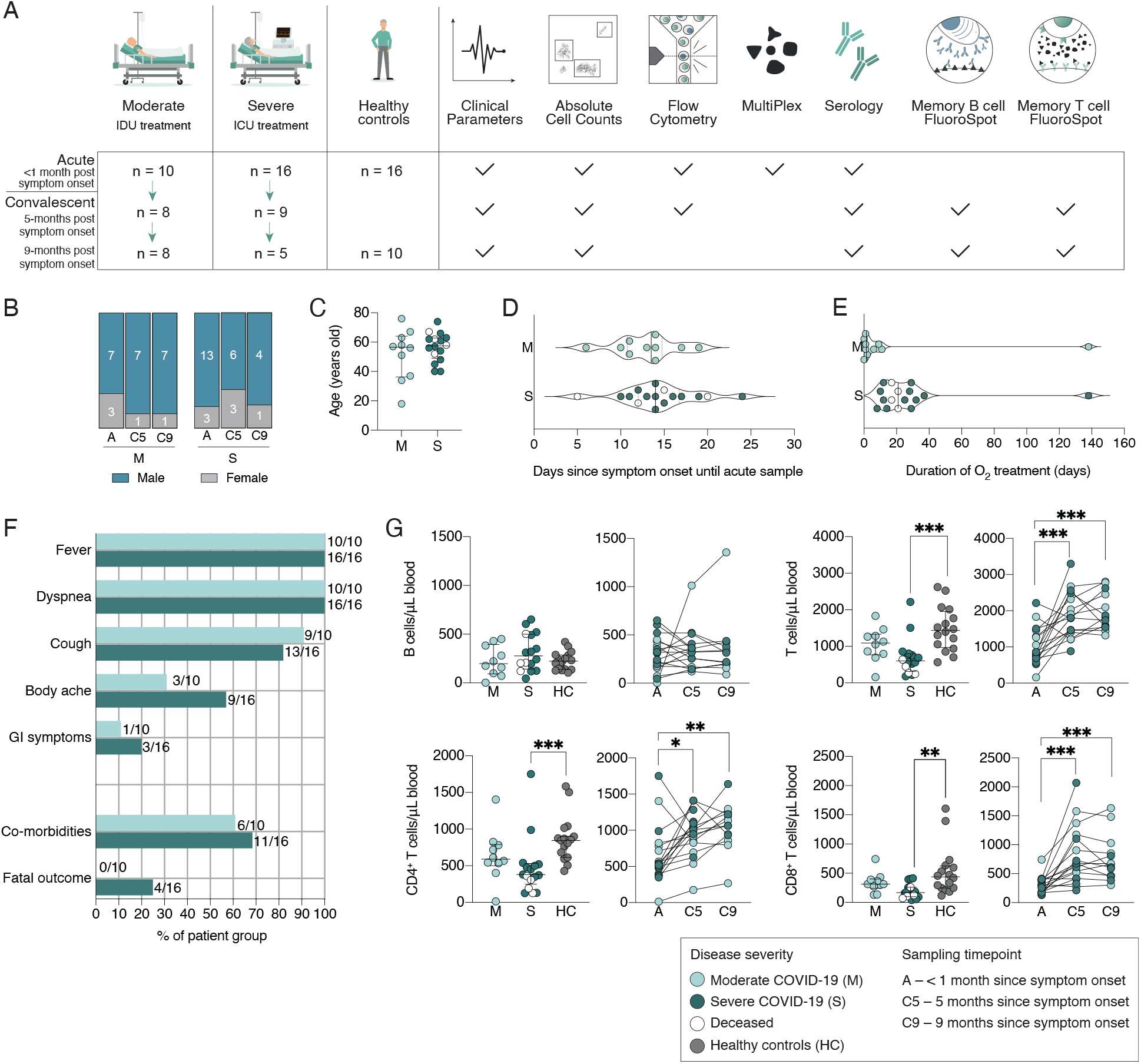
Study design and clinical features of hospitalized COVID-19 patients. (A) Schematic overview of patient cohorts and experimental setup. IDU – infectious disease unit; ICU – intensive care unit. (B) Sex distribution among moderate and severe patients at acute phase, and convalescence. (C) Age distribution among COVID-19 patients. (D) The number of days from symptom onset to acute sampling of COVID-19 patients. (E) The duration of supplemental oxygen treatment among COVID-19 patients during the acute phase. (F) Symptoms, co-morbidities and outcome of COVID-19 patients. GI – gastrointestinal symptoms. (G) Absolute cell numbers in peripheral blood of COVID-19 patients during the acute phase and in healthy controls, as well as longitudinally sampled patients. All scatter plots show median and IQR where n = 10 for moderate, n = 16 for severe, n = 16 for healthy control, n = 17 for C5 and n = 13 for C9. Statistical significance in G (M, S and HC) was assessed by non-paired Kruskal-Wallis test corrected for with Dunn’s multiple comparisons test. Statistical significance in longitudinal plots of G (A, C5 and C9) was assessed with Wilcoxon matched pairs signed rank test. *p < 0.05; **p < 0.01; and ***p < 0.001.

**Table 1.** Demographics and clinical characteristics of COVID-19 patients.-1.

|  | Acute COVID-19 |  | Convalescent COVID-19<br>5 months |  | Convalescent COVID-19<br>9 months |  |
| --- | --- | --- | --- | --- | --- | --- |
|  | Moderate | Severe | Moderate | Severe | Moderate | Severe |
| Total n | 10 | 16 | 8 | 9 | 8 | 5 |
| Age, years, median (range) | 57 (18-76) | 58 (40-74) | 58 (34-76) | 56 (40-63) | 58 (34-76) | 61 (45-63) |
| Male, n | 7 | 13 | 7 | 6 | 7 | 4 |
| Female, n | 3 | 3 | 1 | 3 | 1 | 1 |
| Symptom onset to sampling, days, median (range) | 14 (6-19) | 14 (5-24) | 155 (144-171) | 155 (142-172) | 274 (264-287) | 278 (263-284) |
| Total duration of hospitalization, median days (range) | 8 (5-39) | 19 (10-138) | 8 (5-39) | 17 (10-57) | 8 (5-39) | 15 (10-49) |
| At least one co-morbidity<sup>a</sup>, n | 6 | 11 | 6 | 6 | 6 | 4 |
| Fatal outcome, n | 0 | 4 | 0 | 0 | 0 | 0 |
| **Peak supplemental oxygen treatment** |  |  |  |  |  |  |
| No oxygen, n | 2 | 0 | 1 | 0 | 1 | 0 |
| Low flow, n | 7 | 3 | 6 | 3 | 6 | 2 |
| High flow, n | 1 | 0 | 1 | 0 | 1 | 0 |
| Mechanical ventilation/ECMO, n | 0 | 13 | 0 | 6 | 0 | 3 |
| **Treatment** |  |  |  |  |  |  |
| ICU treatment, n | 0 | 16 | 0 | 9 | 0 | 5 |
| Steroids before acute sampling, n | 2 | 11 | 2 | 5 | 2 | 3 |
| Antibiotics before acute sampling, n | 3 | 11 | 2 | 6 | 2 | 3 |
| Anticoagulants before acute sampling, n | 10 | 14 | 8 | 9 | 8 | 5 |
| Specific treatment before acute sampling<sup>b,c</sup>, n | 0 | 2<sup>b,c</sup> | 0 | 1<sup>b</sup> | 0 | 1<sup>b</sup> |

Abbreviations: ECMO - extracorporeal membrane oxygenation; ICU – intensive care unit.
Footnotes: <sup>a</sup> Hypertension, type II diabetes, asthma, obesity, obstructive sleep apnea syndrome, chronic hepatitis B, coronary heart disease; <sup>b</sup> Remdesivir; <sup>c</sup> Tocilizumab.

Absolute numbers of lymphocyte subsets in peripheral blood were also assessed during the acute phase and at 5- and 9-month convalescence. During the acute phase, there was a significant decrease in total T cell numbers, including CD4^+^ T cell and CD8^+^ T cell subset numbers in the severe group, but not in moderate COVID-19 patients as compared to healthy controls, previously described by Sekine *et al*.^16^ (Figure 1G). Total B cell numbers in peripheral blood of COVID-19 patients were comparable to healthy control levels throughout the study. However, absolute T cell numbers returned to healthy control levels by the 5-month convalescence time point (Figure 1G).

### Increased germinal center activity in COVID-19 patients

Germinal center activity was assessed using the surrogate marker, CXCL13, in hospitalized COVID-19 patients and healthy controls. Levels of plasma CXCL13 were significantly elevated in both moderate and severe patients during acute COVID-19 compared to healthy controls (Figure 2A). Although a trend towards higher CXCL13 levels was observed in the severe patient group compared to moderate patients, this was not statistically significant. This led us to measure the frequencies of activated cTfh cells in peripheral blood as an additional indication of germinal center activity (Figure 2B). A marked increase in the frequencies of total activated cTfh cells (CD4^+^ CXCR5^+^ ICOS^+^ PD1^+^) was observed in both moderate and severe patient groups compared to healthy controls (Figure 2C). Increased activation of Th1-polarized cTfh (CD4^+^ CXCR5^+^ CXCR3^+^ ICOS^+^ PD1^+^) cells was detected in both moderate and severe patients, while the Th2/Th17-polarized subset (CD4^+^ CXCR5^+^ CXCR3^-^ICOS^+^ PD1^+^) was activated only in severe patients (Figure 2D-E). At 5-months post infection, the frequencies of activated cTfh cells declined to normal levels, comparable to healthy controls (Figure 2C-E).

**Figure 2.**
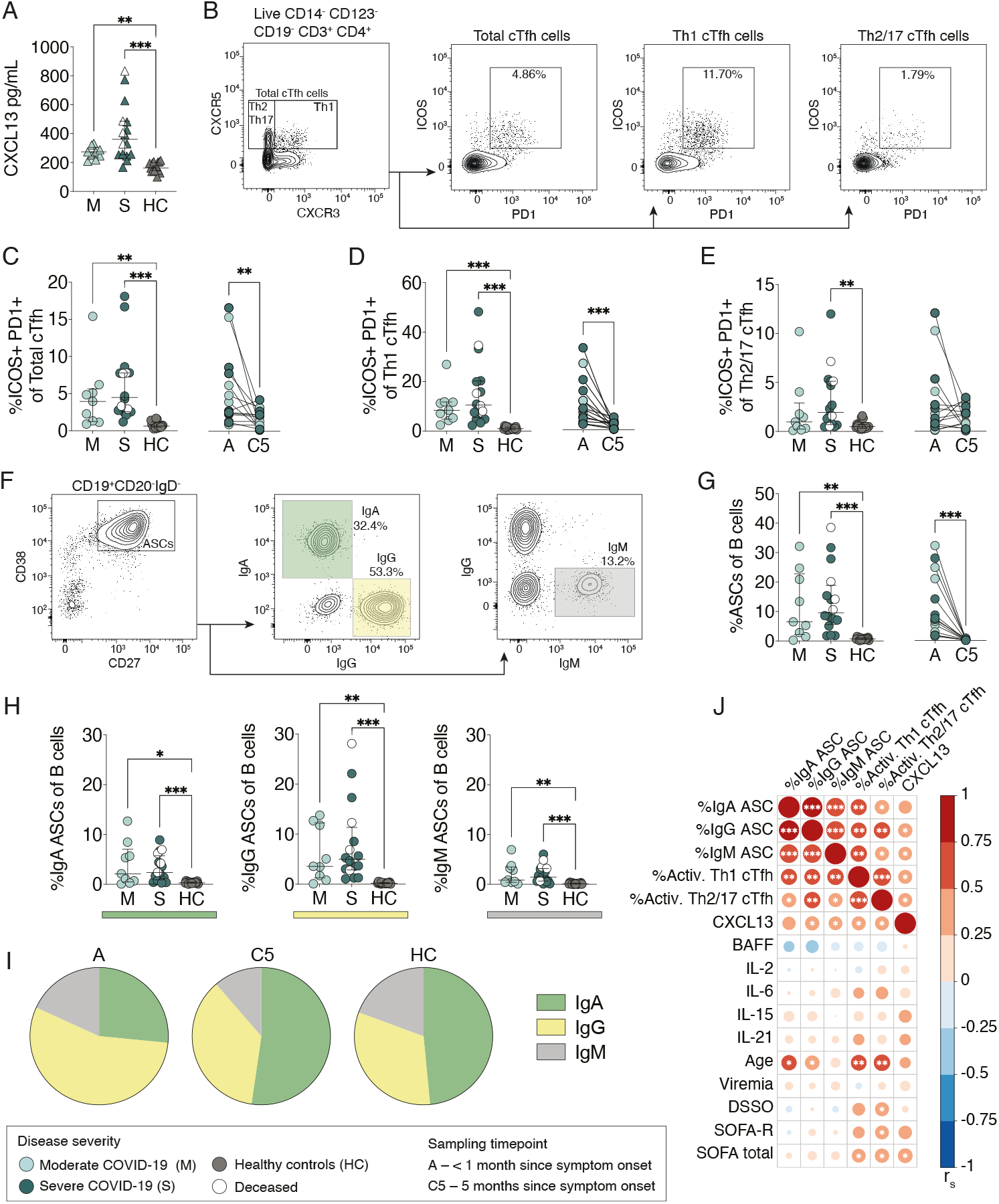
Increased germinal center activity and expansion of antibody-secreting cells in COVID-19 patients. (A) CXCL13 plasma concentrations in moderate and severe COVID-19 patients at the acute phase, as well as in healthy controls. (B) Flow cytometry gating strategy of total, Th1- and Th2/17-polarized cTfh cells in a representative COVID-19 patient. (C-E) Frequencies of activated (ICOS^+^ PD1^+^) total, Th1-polarized, and Th2/17-polarized cTfh cells in COVID-19 patients and healthy controls. Comparisons between moderate and severe patients with healthy controls, as well as longitudinally sampled patients. (F) Flow cytometry gating strategy of antibody-secreting cells (ASCs) and their immunoglobulin expression in a representative COVID-19 patient. (G) Frequencies of ASCs within the total B cell pool in COVID-19 patients and healthy controls. Comparisons between moderate and severe patients with healthy controls, as well as longitudinally sampled patients. (H) Frequencies of IgA-, IgG-and IgM-ASC subsets within the total B cell pool. (I) Distribution of Ig isotype expression by ASCs in COVID-19 patients and healthy controls. (J) Spearman’s correlation matrix where color scale and size of the circles indicate Spearman’s correlation coefficient (r_s_). Data is from COVID-19 patients sampled during the acute phase. DSSO – days since symptom onset; SOFA-R - respiratory sequential organ failure assessment score; SOFA total – total sequential organ failure assessment score. All scatter plots show median and IQR where n = 10 for moderate, n = 16 for severe, n = 16 for healthy control, and n = 17 for C5. Statistical significance in A-H (M, S and HC) was assessed by non-paired Kruskal-Wallis test corrected for with Dunn’s multiple comparisons test. Statistical significance in longitudinal plots of C-G (A and C5) was assessed with Wilcoxon matched pairs signed rank test. *p < 0.05; **p < 0.01; and ***p < 0.001.

### Significant expansion of ASCs dominated by the IgG subset in COVID-19 patients

Freshly isolated PBMCs were analyzed using flow cytometry to determine the magnitude of ASC expansion during acute SARS-CoV-2 infection and at 5-months convalescence (Figure 2F). A significant increase in ASC frequencies was observed in both moderate and severe patients during the acute phase compared to healthy controls (Figure 2G). Interestingly, severe patients who later deceased showed an above median frequency of total ASCs (Figure 2G). At 5-months after the onset of symptoms, ASC frequencies in COVID-19 patients decreased to healthy control levels (Figure 2G). Acute ASC expansion was further assessed for surface and intracellular expression of IgA, IgG and IgM (Figure 2F). A significant expansion in all three Ig-ASC subsets was observed in both moderate and severe patients compared to healthy controls (Figure 2H). The ASC expansion was dominated by the IgG-ASC subset whereas IgA-ASCs dominated during convalescence and in healthy controls (Figure 2I). We observed significant associations between ASC expansion, cTfh activation and CXCL13 levels as well as age, but no associations were found between these parameters and viremia, BAFF, IL-2, IL-21, IL-15 or IL-6 plasma levels at acute sampling (Figure 2J). To summarize, an active germinal center response was observed as indicated by increased CXCL13 levels in plasma and the activation of cTfh cells resulting in the expansion of ASCs during acute infection with SARS-CoV-2.

### SARS-CoV-2-specific antibody levels during acute COVID-19 are associated with disease severity

We next assessed the levels and dynamics of SARS-CoV-2-specific antibodies during the acute phase of COVID-19, as well as at 5- and 9-months since symptom onset (n = 26, 17 and 13, respectively). During the acute phase of COVID-19, 23 out of the 26 patients had neutralizing antibody titers of >10 (Figure 3A). The three patients with undetectable neutralizing antibodies were also negative for IgG/IgM specific to SARS-CoV-2 receptor-binding domain (RBD). However, 18 and 21 out of the 26 patients had detectable levels of spike subunit 1 (S1)-specific and nucleocapsid (N)-specific IgG antibodies, respectively, at the acute phase (Figure 3A). A higher proportion of patients negative for S1- and N-IgG were in the moderate COVID-19 patient group, while severe patients had significantly higher SARS-CoV-2-specific and neutralizing antibody levels, despite comparable sampling timepoints between the two patient cohorts (Figure 1D and Figure 3A). Neutralizing antibody titers, as well as S1- and N-specific IgG levels were higher at later timepoints during the acute phase (Figure 3B). All patients sampled later than 15 days after symptom onset had detectable levels of both S1- and N-IgG as well as neutralizing antibodies. The four patients that later deceased had neutralizing antibody titers between 120 and 960. A strong positive correlation between S1-IgG, N-IgG, and neutralizing antibody titers were observed during the acute phase, and both S1- and N-IgG levels correlated with the number of days since symptom onset (Figure 3C). Higher antibody levels during acute COVID-19 were also associated with degree of respiratory failure (SOFA-R), but not age (Figure 3C).

**Figure 3.**
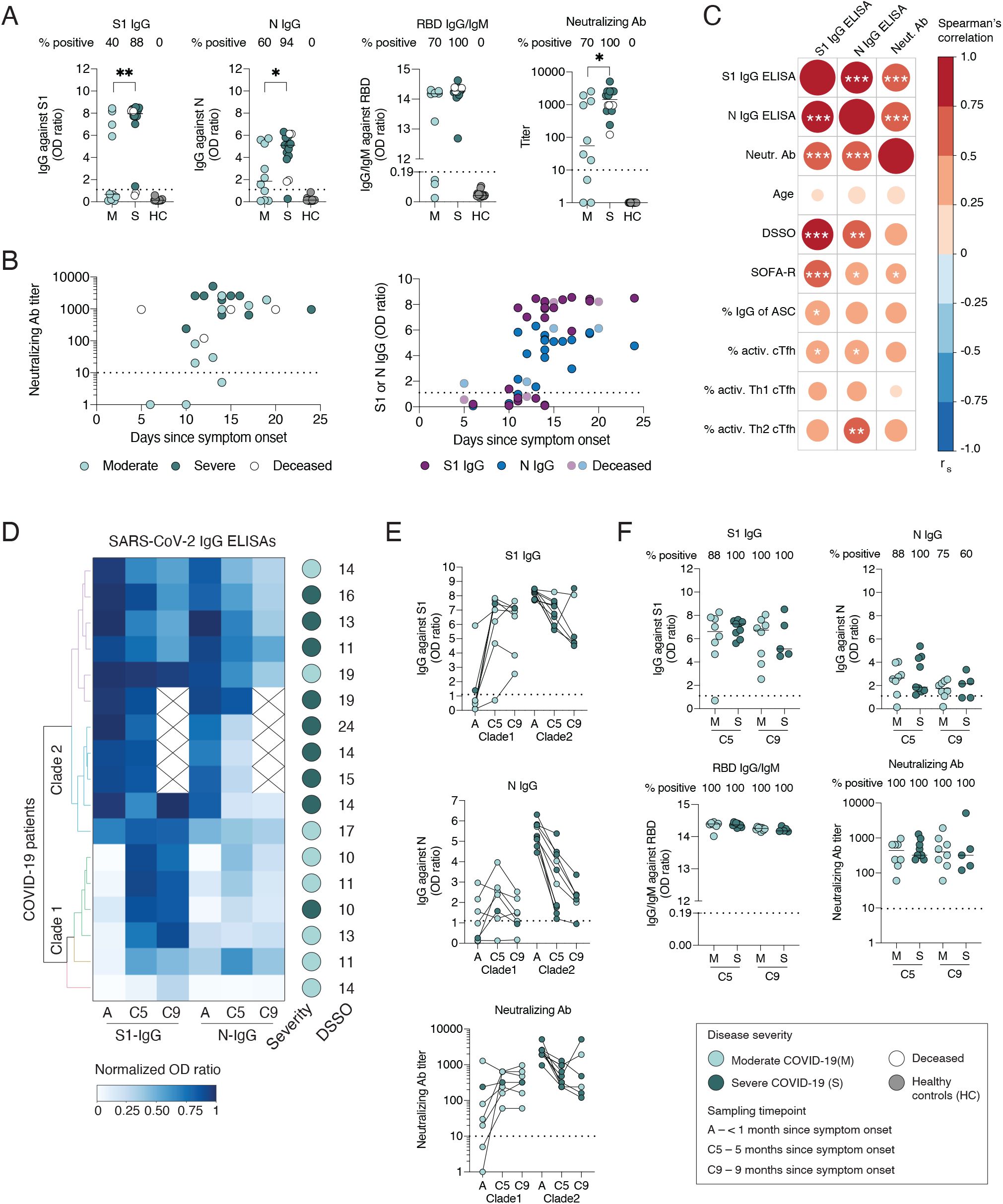
SARS-CoV-2-specific antibody levels and dynamics during acute COVID-19 and in convalescence. (A) S1-IgG and N-IgG antibody levels, positivity for RBD-IgG/IgM, and SARS-CoV-2-neutralizing antibody titers during acute phase of COVID-19 in moderate and severe patients, as well as healthy controls. (B) S1-IgG and N-IgG antibody levels, and neutralizing antibody titers in regard to the number of days since COVID-19 symptom onset. (C) Spearman’s correlation matrix where color scale and size of the circles indicate Spearman’s correlation coefficient (r_s_). Data is from all COVID-19 patients sampled during the acute phase. DSSO – days since symptom onset, ASC – antibody-secreting cells, SOFA-R – respiratory sequential organ failure assessment score. (D) Clustering analysis based on S1-IgG and N-IgG levels in longitudinally followed up COVID-19 patients. The blue color scale indicates normalized OD ratio values from S1-IgG and N-IgG ELISAs. DSSO – days since symptom onset. (E) S1-IgG, N-IgG, and neutralizing antibody levels over time in COVID-19 patients with paired samples divided in two clades based on clustering analysis in D. (F) Comparison of antibody levels between moderate and severe COVID-19 patients at 5- and 9-months after symptom onset. All scatter plots show median where n = 26 for acute phase, n = 17 for C5, n = 13 for C9, and n = 16 for HC. Dotted horizontal lines represent threshold for positivity in each assay (defined by the manufacturer for ELISAs, and <10 for neutralization). Statistical significance was assessed using non-paired Kruskal-Wallis test corrected for with Dunn’s multiple comparisons test. *p < 0.05; **p < 0.01; and ***p < 0.001.

### SARS-CoV-2-specific antibodies persist up to 9-months after symptom onset

Next, we assessed the dynamics of antibody levels in COVID-19 patients sampled longitudinally. Clustering analysis based on S1- and N-IgG antibody levels was performed at the acute phase and at 5- and 9-months since symptom onset (Figure 3D). The patients clustered into two major clades based on the dynamics of their antibody levels (Figure 3D). Patients in clade 1 mainly exhibited increased SARS-CoV-2-specific IgG levels at 5 months as compared to the acute phase, followed by a decrease in antibody levels at 9 months (Figure 3E). Meanwhile, patients in clade 2 exhibited decreased antibody levels at both 5 and 9 months, as compared to the acute phase (Figure 3E). Patients from clade 1 were sampled relatively early in the acute phase (a median of 11 days since symptom onset), but also primarily consisted of moderate patients, while patients in clade 2 were sampled slightly later in the acute phase (median 15 days since symptom onset), and primarily consisted of severe patients. Neutralizing antibody titers in both clades remained relatively unchanged between 5month and 9-month time points. Notably, one patient in the severe group had a neutralizing antibody titer of 1920 at the acute phase, and a titer of 480 at 5 months, but reached a titer of 5120 at 9 months, suggesting a potential SARS-CoV-2 reexposure. S1-IgG, but not N-IgG, levels also increased in this patient at 9 months (Figure 3E).

At 5 and 9 months, all followed up patients had detectable levels of S1-IgG, RBD-IgG/IgM and neutralizing antibodies, while N-IgG levels decreased to borderline positive values in four patients (Figure 3F). Neutralizing antibody titers ranged from 60 to 1860 in moderate and 120 to 5120 in severe patients (Figure 3F). However, there was no significant difference in antibody levels between moderate and severe patients, suggesting that regardless of the level of care required during acute COVID-19, hospitalized patients maintain comparable antibody levels in convalescence (Figure 3F).

### Robust S1- and N-specific B cell memory persists up to 9 months after symptom onset

We next assessed the presence of SARS-CoV-2-specific memory B cells in convalescent COVID-19 patients at 5- and 9-month post symptom onset, and in pre-pandemic healthy controls (Figure 4A and Figure S2). We utilized a FluoroSpot assay which allows detection of S1- and N-specific memory B cell-derived antibody-secreting cells (mASCs), secreting either IgG or IgA following polyclonal B cell stimulation (Figure 4A and Figure S2). S1-specific IgG-mASCs were detected in all patients at both 5 and 9 months, while N-specific IgG-mASCs were detected in all but one patient who was below positive threshold at 9-months following symptom onset (Figure 4B). Very few patients had detectable S1- or N-specific IgA-mASCs after stimulation (Figure S2).

**Figure 4.**
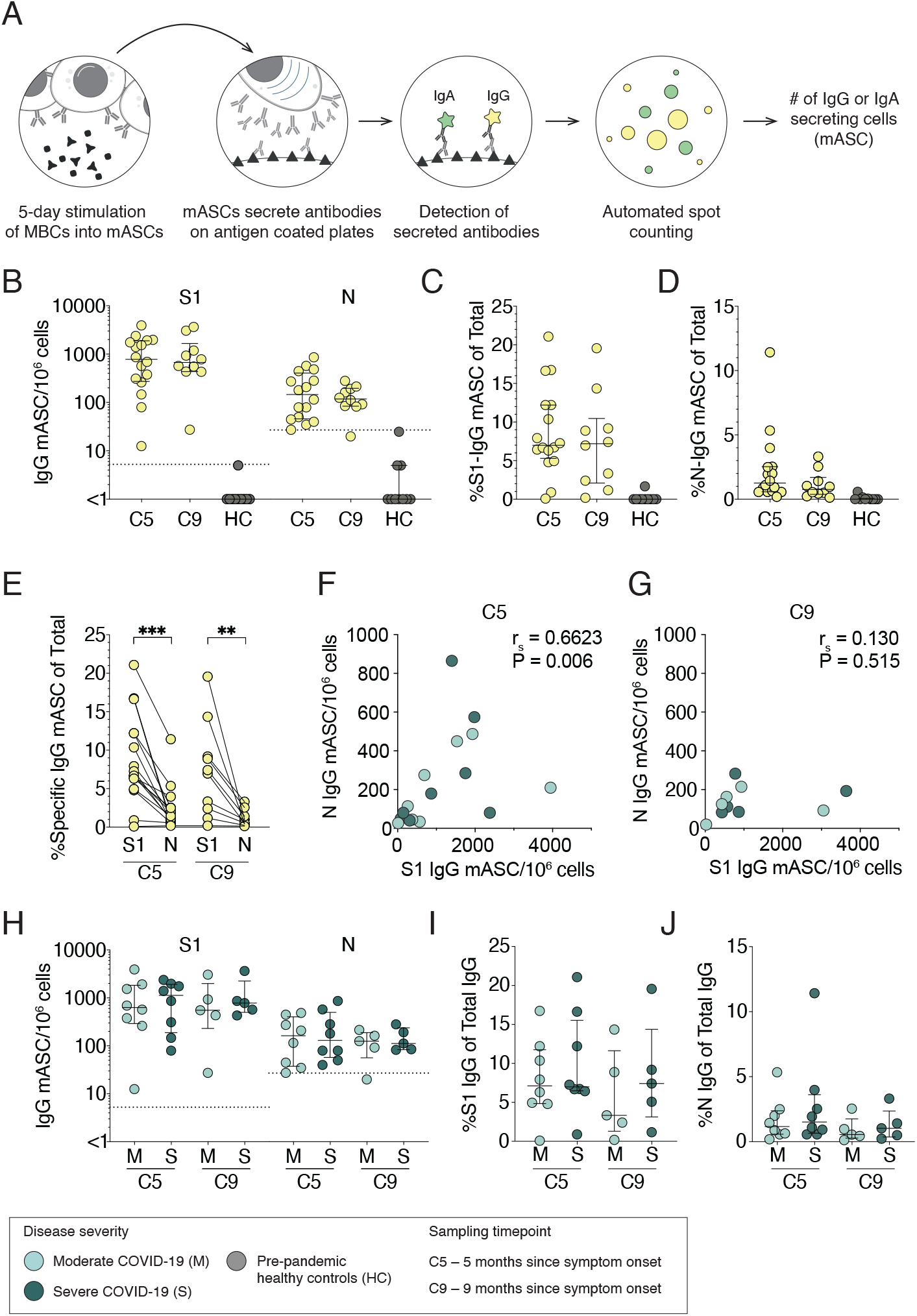
Robust S1- and N-specific B cell memory persists up to 9-months after symptom onset. (A) Schematic of memory B cell FluoroSpot assay. (B) Numbers of S1- and N-specific memory B cell-derived IgG-ASCs (mASCs) at 5- and 9-months in COVID-19 patients and pre-pandemic healthy controls. (C) Frequencies of S1-specific mASCs within total IgG-mASC pool following polyclonal B cell stimulation. (D) Frequencies of N-specific mASCs within total IgG-mASC pool following polyclonal B cell stimulation. (E) Comparison between the frequencies of S1-specific and N-specific IgG mASCs within total mASC pool for each COVID-19 patient at 5- and 9-months. (F and G) Spearman correlation between S1-specific and N-specific mASCs numbers after stimulation in COVID-19 patients at 5- and 9-months. rs - Spearman’s correlation coefficient. (H-J) Comparison between moderate and severe COVID-19 patients in regard to numbers and frequencies of S1- and N-specific mASC at 5- and 9-months. Dotted threshold lines in (B) and (H) defined by average S1- or N-mASCs numbers in pre-pandemic HCs plus three standard deviations. Median with IQR is plotted in all scatter plots where n = 16 for C5, n = 10 for C9, n = 10 for HC. Statistical significance in E was assessed by Wilcoxon matched-pairs signed rank test. **p < 0.01; and ***p < 0.001.

To determine the relative abundance of SARS-CoV-2-specific memory B cells within each patient, we measured the frequencies of S1- and N-specific mASC numbers within the total IgG-secreting mASC pool after the stimulation. A median of 7% out of the total IgG-mASC were S1-specific, while a median of 1% out of the total IgG-mASC were N-specific in COVID-19 patients (Figure 4D). However, there was a high variation in frequencies of S1- or N-specific cells between patients, ranging from 0.2% to 21% (Figure 4C and D). Consistently higher frequencies of S1-specific IgG-mASCs were observed after stimulation, as compared to N-specific IgG-mASCs (Figure 4E). The numbers of S1-specific mASCs were shown to positively correlate with the numbers of N-specific mASCs at 5 months post symptom onset, but not at 9 months (Figure 4F and 4G). Finally, disease severity during the acute phase of COVID-19 had no effect on the magnitude of memory B cell response in convalescence (Figure 4H-J).

### Polyfunctional SARS-CoV-2-specific T cell memory persists up to 9 months after symptom onset

To determine if T cell memory is generated and sustained in COVID-19 patients we utilized a FluoroSpot assay to simultaneously measure the number of T cells secreting IFN-γ, IL-2 and/or TNF in response to stimulation with SARS-CoV-2 peptides (Figure 5A; Figure S2). In addition to determining SARS-CoV-2-specific T cell numbers, relative amounts of secreted cytokines, as well as cytokine co-expression patterns were assessed, allowing for a characterization of polyfunctional T cell responses. PBMCs from COVID-19 patients sampled at 5- and 9-months since symptom onset as well as pre-pandemic healthy controls were stimulated with three different SARS-CoV-2 peptide pools (Figure 5B). Numbers of all cells secreting cytokines after peptide stimulation were first enumerated by detecting either IFN-γ, IL-2 or TNF (Figure 5C). Total IFN-γ and IL-2-secreting cell numbers, but not TNF-secreting cell numbers, were higher in COVID-19 patients as compared to healthy controls. High numbers of TNF-only producing cells in healthy controls may be due to unspecific TNF-secretion by other PBMC subset than T cells. Next, we assessed relative amounts of IFN-γ, IL-2 or TNF secreted by responding cells. Polyfunctional cells, especially those secreting all three cytokines at once, on average secreted more cytokines than single-functional cells (Figure 5D). This led us to investigate cytokine co-expression patterns and determine the numbers of polyfunctional cells responding to SARS-CoV-2 peptide pools. A large fraction of cells responding to SARS-CoV-2-peptide stimulation were indeed polyfunctional T cells, with comparable co-expression patterns at 5 months and 9 months after infection (Figure 5E). Robust polyfunctional T cell responses to at least one of the SARS-CoV-2 peptide pools were observed at 5 or 9 months (Figure 5F). Interestingly, we observed significantly more polyfunctional T cells responding to the S1 peptide pool as compared to the N peptide pool, highlighting the immunogenicity of S1 (Figure 5G). Patients with higher S1-specific polyfunctional T cell numbers also had higher N-specific polyfunctional T cell numbers, although this correlation was not significant at the 9-month timepoint (Figure 5H). We did not observe significant differences between moderate and severe COVID-19 patients in regard to the number of polyfunctional T cell responses, at either 5- or 9-months after infection (Figure 5I).

**Figure 5.**
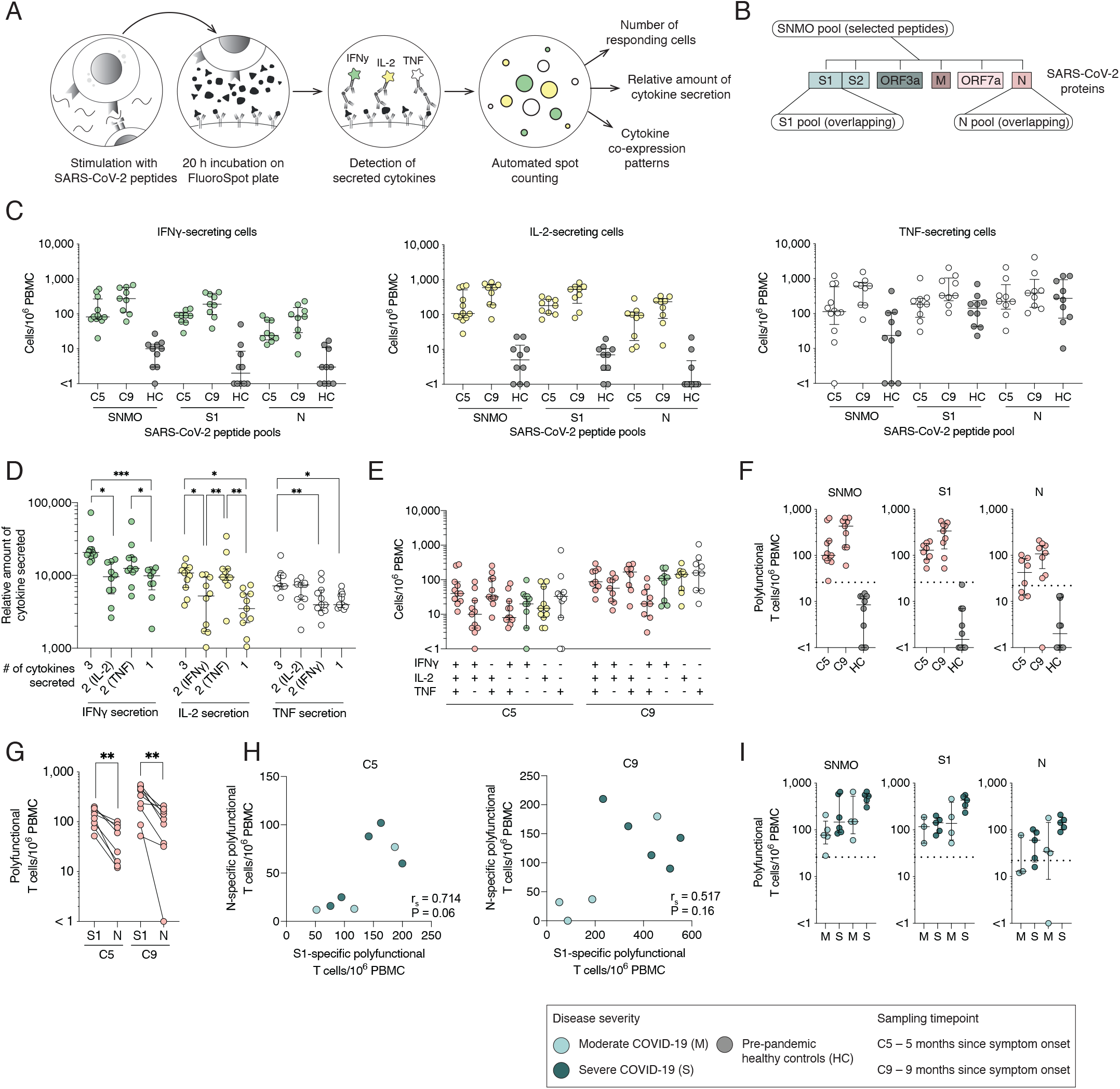
Polyfunctional SARS-CoV-2-specific T cell memory persists up to 9 months after symptom onset. (A) Schematic of memory T cell FluoroSpot assay. (B) SARS-CoV-2 peptide pools used in memory T cell stimulation. (C) Total number of cells responding to stimulation by secreting IFN-γ, IL-2 or TNF at 5- and 9-month convalescence and in pre-pandemic healthy controls. (D) Average spot volume (relative amount of cytokine secreted) for IFN-γ, IL-2 or TNF produced by cells secreting all three, two or one cytokine after SNMO peptide pool stimulation at 5-months. (E) Cytokine co-expression patterns of responding T cells after stimulation with SNMO peptide pool at 5- and 9-months. (F) Total number of polyfunctional T cells (cells secreting at least two cytokines) after stimulation. (G) Comparison between the numbers of polyfunctional T cells after the stimulation with S1 or N peptide pools for each patient. (H) Spearman correlation between S1- and N-specific polyfunctional T cell numbers at 5- and 9-months. rs – Spearman’s correlation coefficient. (I) Comparison of polyfunctional T cell numbers responding to SARS-CoV-2 peptide pools between moderate and severe patients at 5- and 9-months. Median and IQR is plotted in all scatter plots where at 5-months n = 11 for SNMO, n = 8 for S1, n = 8 for N; and at 9-months n = 9 for SNMO, n =9 for S1, n = 9 for N; and for HC n = 10 for SNMO, S1 and N. Dotted horizontal lines represent threshold for positive response defined by average polyfunctional T cell numbers in pre-pandemic HCs after stimulation plus three standard deviations. Statistical significance was assessed using paired non-parametric Friedman test in D or Wilcoxon test in G. *p < 0.05; **p < 0.01; and ***p < 0.001.

## Discussion

The COVID-19 pandemic is one of the largest global health challenges we have faced in the last century. It is hence imperative to assess the development and the longevity of adaptive immunity in infected persons. In this study, we systematically assessed the initiation and longevity of specific humoral and cellular immune responses to SARS-CoV-2 in moderately and severely sick COVID-19 patients, who were longitudinally sampled at the acute phase and at 5- and 9-months after symptom onset.

The assessment of germinal center activity during acute COVID-19 provides insights into the initiation of the adaptive immunity, specifically humoral responses. The activation of germinal centers give rise to both short- and long-lived ASCs as well as memory B cells that will respond quickly upon reexposure to SARS-CoV-2. The absence of germinal centers was recently observed in lymph nodes and spleens from *post mortem* COVID-19 patients^23^ suggesting that SARS-CoV-2 infection, in some cases, may not trigger efficient development of protective immunity and result in fatal outcome. Although we did not assess germinal center activity directly within lymph nodes, we used plasma CXCL13 levels as a surrogate marker^4^, and found increased levels of CXCL13 in both moderate and severe patients. Other recent studies have also observed increased levels of CXCL13 with higher levels in severe, compared with non-severe COVID-19 patients^24^. In the present patient cohorts, there was no significant difference in CXCL13 levels between moderate and severe COVID-19, however, a few patients in the severe group had distinctly high levels. In addition, we observed a significant activation of antiviral Th1-polarized cTfh cells, another indication of germinal center activity, in both patient groups, a result consistent with other studies^16,25–30^. We also found that higher levels of CXCL13 in plasma positively correlated with the activation of cTfh cells. Our findings suggest that germinal center reactions have indeed taken place in secondary lymphoid organs during acute COVID-19 potentially leading to ASC expansion as well as contribution to the generation of long-lived plasma cells and memory B cells.

The expansion of ASCs in peripheral blood has been well documented during acute COVID-19 disease and shown to be SARS-CoV-2-specific^25,31,32^. We also observed a characteristic increase in ASC expansion during acute COVID-19 in both severe and moderate patients dominated by the IgG subset, however no difference was observed between the two patient groups. Expansion of ASCs during viral infections have been shown to be a good predictor of the development of neutralizing antibodies and B cell memory^10^. Additionally, during viral infections, ASCs can produce large amounts of antibodies as long as viral shedding occurs, suggesting that ASCs play an active role in infection clearance^11^. We found that the majority of patients had neutralizing antibody titers during the acute phase, perhaps originating from the expanded ASC population. Taken together, the observed ASC expansion during acute COVID-19 may play an important role in SARS-CoV-2 clearance and disease control.

A major function of humoral responses during acute infection is to generate memory B cells and long-lived plasma cells that will produce pathogen-specific antibodies. Numerous studies have characterized seroconversion kinetics during the acute phase of COVID-19^33–35^. Although the majority of patients seroconvert between 7 to 14 days after symptom onset, a large variation in SARS-CoV-2-specific antibody levels has been observed^34,36–39^. Therefore, sampling timepoint during acute COVID-19 may determine the antibody levels detected within each patient, and patients sampled earlier may have lower antibody levels as compared to sampling later during the acute phase of infection. In this study, we confirm that seropositivity for SARS-CoV-2 antibodies is time-dependent, as higher antibody levels are observed in patients sampled later during the acute phase. However, moderate and severe COVID-19 patients in this study were sampled at comparable timepoints during the acute phase, yet severe patients had significantly higher SARS-CoV-2-specific antibody levels compared to moderate patients. Our results confirm previous reports^30,40^, showing a positive relationship between degree of disease severity and antibody levels during acute COVID-19.

Longevity of immunological memory following SARS-CoV-2 infection is a question of particular relevance for informing public health policies and predicting the future course of the pandemic. Several studies have demonstrated decreasing antibody levels in COVID-19 patients over time, leading to concerns for waning immunity^41–43^. However, for a more comprehensive picture of the longevity of immunity, memory B cell and T cell responses should be studied in parallel, as these cells would be important players in providing protection upon re-exposure to SARS-CoV-2. Here, we followed 17 patients for 5-months, and 13 of those patients for 9-months after symptom onset, in order to assess antibody dynamics over time. Consistent with other studies, we show a substantial decline in antibody levels over time, however, all patients followed up to 5-months and 9-months maintain detectable neutralizing antibody titers^12,13^. In parallel, we measured memory B cell and also T cell responses in the same individuals utilizing FluoroSpot assays. Recent studies have shown that memory B cells and T cells were detectable up to 8 months post infection^12,13^. Here, we confirm these observations as S1- and N-specific IgG memory B cells as well as specific-memory T cells are readily detectable in circulation at both 5- and even 9-months post symptom onset in all patients within this cohort, although magnitude of these responses is highly variable. We consistently observed a lower magnitude of N-specific B cell and T cell memory responses, as compared to S1-specific responses. These results confirm the previously demonstrated preferential targeting of spike protein over nucleocapsid protein^12,16,29,31,44^.

Functional T cell responses to SARS-CoV-2 have been detected in both the acute phase of disease and in convalescence by numerous studies^12,16,30,44,45^. In other disease settings, it has been shown that polyfunctional T cells (secreting more than one cytokine after stimulation) are important contributors to the control of viral infections^46–49^. These polyfunctional T cells secrete higher amounts of cytokines on a cell basis, as compared to single-functional T cells^49–51^. In this study, polyfunctional T cell responses to SARS-CoV-2 peptides were detectable in all COVID-19 patients, although at varying magnitude. Polyfunctional T cells, especially cells co-secreting all three cytokines measured (i.e., IFN-γ, IL-2 and TNF), were also shown to, on average, produce higher amounts of each cytokine than single-functional cells. We highlight the strength of analyzing secretion of multiple cytokines at once, as this approach provides information on cytokine co-expression patterns and identification of polyfunctional T cells, which are superior in their cytokine secretion capacity and therefore may be important in antiviral defense upon re-exposure to SARS-CoV-2.

The role of adaptive immunity and protection from clinical disease after re-exposure to SARS-CoV-2 is under active investigation. The exact magnitude of adaptive immune responses to SARS-CoV-2 infection required for protection from reinfection remains unclear. However, recent longitudinal studies on larger cohorts suggest that the presence of cellular and humoral immunity is highly associated with prevention from re-infection with SARS-CoV-2 and clinical disease^52–54^. Although we did not follow up patients in this cohort regarding PCR-positivity for SARS-CoV-2 after the acute phase, none of the patients developed clinical COVID-19 for a second time since recovery and throughout the course of this study.

The differences in antibody levels during the acute phase of disease between moderate and severe COVID-19 patients were not apparent during late convalescence. Neither neutralizing antibody titers, nor the magnitude of B cell and T cell memory responses differed between moderate and severe patients during late convalescence at either 5- or 9-months. This suggests that regardless of the disease severity and level of care required during the acute phase, the majority of hospitalized patients are likely to elicit a long-lasting cellular and humoral immunity towards SARS-CoV-2.

In summary, we demonstrate the activation of germinal centers, expansion of ASCs and the presence of SARS-CoV-2-specific antibodies in both moderate and severe patients during acute COVID-19. SARS-CoV-2 specific and neutralizing antibodies gradually declined after the acute phase in most patients but were sustained through convalescence. SARS-CoV-2-specific memory B cells and T cells, assessed at 5- and 9-months symptom onset were also detectable in peripheral blood in all COVID-19 patients, but at heterogenous magnitude, and a strong polyfunctional memory T cell response was observed. Taken together, the present findings reveal that hospitalized patients develop and sustain robust SARS-CoV-2-specific adaptive immune responses that last up to at least 9-months post symptom onset.

### Limitations of Study

The current study, although extensive in regard to immunological analyses performed, is limited by a relatively small COVID-19 patient cohort size (acute phase n = 26, 5-months n = 17, and 9-months n = 13). Here, we assessed SARS-CoV-2-specific immune responses in moderately and severely sick COVID-19 patients who required hospitalization, however, we were not able to include asymptomatic or mild cases. Therefore, the longitudinal characteristics and magnitude of the immune responses observed in the current COVID-19 patient cohort may only be applicable to hospitalized patients.

## Materials & Methods

### Ethics statement

The study was approved by the Swedish Ethical Review Authority. All patients or next of kin and control donors provided informed consent in line with the ethical approval.

### Study subjects and sampling

Twenty-six hospitalized SARS-CoV-2 PCR-confirmed patients from Karolinska University Hospital in Stockholm, Sweden, were included in the study in April and May 2020. Moderate COVID-19 patients were recruited at the infectious disease unit (IDU), while severe COVID-19 patients were recruited at the intensive care unit (ICU). For supplemental oxygen treatment requirement and additional treatments during the acute phase for each group see Table 1. Sequential organ failure assessment (SOFA total) score, including respiratory SOFA (SOFA-R) score, was also used to describe the severity of COVID-19 disease^55^. Patients were primarily male, with median age of 57 (range 18-76) for moderate and median age of 58 (range 40-74) for severe patients (Figure 1A-C). Patients were sampled from 5 to 24 days since symptom onset during the acute phase (Figure 1D). In addition, 17 patients were followed up at 5months and 13 patients were followed up at 9-months since symptom onset (Figure 1A). This COVID-19 patient cohort is part of the Karolinska KI/K COVID-19 Immune Atlas project^16–21^. More clinical information and other related data can be found at covid19cellatlas.com. Sixteen healthy age and sex-matched SARS-CoV-2 seronegative controls from 2020 were included for flow cytometry, multiplex and serology experiments (Figure 1A). An additional ten pre-COVID-19 pandemic controls from 2017 (age range 18-50 years) were included for memory B cell and T cell FluoroSpot experiments only. Venous blood samples from study participants were collected in serum and heparin tubes. PBMCs were isolated using gradient centrifugation and used for fresh experiments or cryopreserved. Serum tubes were allowed to stand upright for 2 hours at room temperature (RT) and serum was isolated by centrifugation at 2000 X *g* for 10 minutes and stored at −80°C for later analysis.

### Absolute cell counts

Absolute numbers of B cells and T cells in peripheral blood were measured using BD Multitest™ 6-color TBNK reagents and BD Trucount™ Tubes (BD Biosciences) according to the manufacturer’s instructions. Briefly, exactly 50 *μ*1 of heparinized anti-coagulated blood was added to Trucount tubes within 3 hours after extraction and stained for 15 minutes at RT in the dark. Samples were then fixed with 1X BD FACS lysing solution before acquiring data on a FACSSymphony A5 flow cytometer (BD Biosciences).

### Flow cytometry

Freshly isolated PBMCs from patient and healthy control blood samples were stained using the following reagents and antibodies: Live/Dead cell marker Aqua (Life Sciences), anti-CD19 BUV 395, anti-CD4 Qdot 605, anti-CD38 Pacific Blue, anti-CD14 AmCyan, anti-Ki67 AF700, anti-CD20 FITC, anti-CD123 AmCyan, anti-CD27 BV650, anti-IgD PE-Cy7, anti-IgG PE, anti-CD3 PE-Cy5, anti-IgA APC, anti-IgM BV785, anti-ICOS APC-Cy7, anti-CXCR5 BV711, anti-CXCR3 PE-Dazzle, anti-PD1 BUV737 (Biolegend, BD, Beckman Coulter). Briefly, cells were incubated with 50 μl of surface staining antibody mix diluted in PBS for 30 minutes at 4°C in the dark. Following incubation, cells were washed twice in FACS buffer (2% FCS and 2 mM EDTA in PBS) and then fixed and permeabilized using Foxp3/Transcription Factor Staining kit (eBioscience) for 30 minutes at 4°C. Cells were then washed twice in permeabilization buffer (eBioscience) followed by intracellular staining. Antibodies against Ki67, IgG, IgA and IgM in permeabilization buffer were added to the cells and incubated in the dark for 30 minutes at 4°C. Cells were washed with FACS buffer and then fixed for 2 hours in 1% PFA. Samples were acquired with a BD LSRFortessa (BD Biosciences) followed by analysis with FlowJo software version 10 (FlowJo Inc). For flow cytometry gating strategy see Figure S3.

### SARS-CoV-2-specific antibody ELISAs

RBD-specific IgG/IgM antibodies was assessed using WANTAI SARS-CoV-2 Ab ELISA (Beijing Wantai Biological Pharmacy Ent.), while S1- and N-specific IgG antibody levels were assessed using semi-quantitative IgG ELISAs (Euroimmun) according to the manufacturer’s instructions.

### Microneutralization assay

The neutralizing capacity of antibodies against SARS-CoV-2 in patient serum was tested as previously described^56^. Briefly, heat-inactivated serum samples (30 minutes at 56 °C) were diluted in two-fold dilution series from 1:5 to 1:5120 in Eagle’s Minimum Essential Medium (Gibco) with 5% FCS (Thermo Fisher Scientific). The dilutions were subsequently mixed with equal volumes of 4000 TCID50/ml SARS-CoV-2 (200 TCID50/ well) resulting in final serum dilution series from 1:10 to 1:10240. Final serum dilutions were incubated for 1 hour at 37°C and 5% CO_2_ in duplicate and then moved to 96-well plates seeded with confluent Vero E6 cells, followed by a 4-day incubation at 37°C 5% CO_2_. At the end of incubation, the cells were inspected via optical microscopy for signs of cytopathic effect (CPE). Samples were considered neutralizing if less than 50% of the cell layer showed signs of CPE and nonneutralizing if ≥50% CPE was observed. Results are given as the arithmetic mean of the reciprocals of the highest neutralizing dilutions of the duplicates from each sample.

### rSARS-CoV-2 N protein

A recombinant SARS-CoV-2 N protein was produced by transforming a synthesized plasmid coding for N protein into *Escherichia coli* as previously described^31^. Purified protein was later used in B cell FluoroSpot assays.

### Memory B cell FluoroSpot assay

Detection of SARS-CoV-2 spike subunit 1 (S1) and nucleocapsid (N) protein-specific IgA and IgG memory B cell derived ASCs (mASCs), as well as the total number of IgA- and IgG-mASCs, were measured using a multicolor B cell FluoroSpot kit with modifications (Mabtech). Briefly, thawed, cryopreserved PBMCs from study subjects were stimulated for 5 days with polyclonal B cell stimulation (1μg/mL R848 and 10 ng/mL rIL-2, Mabtech) in R10 media (RPMI 1640 with 10% FCS, 1% penicillin/ streptomycin, and 2 mM L-glutamine, Thermo Fisher Scientific) at 37°C in 5% CO_2_ to differentiate quiescent MBCs into mASCs. Ethanol-activated low autofluorescent polyvinylidene difluoride membrane plates were coated overnight with either 1) anti-IgA and anti-IgG capture antibodies (15 mg/mL) or 2) SARS-CoV-2 S1 (20 μg/ mL) or N protein (10 mg/mL). Excess coating solutions were removed, and plates were washed with PBS before being blocked with R10 for 30 minutes at RT. Optimized dilutions of 5-day stimulated cells were carefully transferred to the coated plates and incubated at 37°C in 5% CO_2_ for 20 hours. Cells were then discarded from plates and the captured antibodies were developed with anti-human IgG-550 and anti-human IgA-490 detection antibodies (1:500 dilution). Fluorescent spots indicating a single mASC and relative volume of antibody produced were detected with an IRIS FluoroSpot Reader and counted with Apex software (Mabtech). Positivity thresholds were set for both S1 and N mASCs based on prepandemic control samples’ average S1 or N plus three standard deviations.

### SARS-CoV-2 peptides for T cell FluoroSpot

Three peptide pools were used in memory T cell FluoroSpot assay: (i) SNMO defined peptide pool (Mabtech) consisting of 47 peptides (8 – 18 amino acid long) from spike (S), nucleocapsid (N), membrane (M), open reading frame (ORF)-3c and ORF-7a proteins, with purity of 60-99%; (ii) S1-spanning overlapping peptide pool (Miltenyi) of 15-mer sequences and 11 amino acid overlap with purify of > 70%; and (iii) N-spanning overlapping peptide pool (Miltenyi) of 15-mer sequences and 11 amino acid overlap with purify of > 70%. Lyophilized peptide pools were reconstituted in 20% DMSO solution at a stock concentration of 200 μg mL for SNMO, and 50 μg/mL for both S1 and N and later diluted to 2 μg/mL for the FluoroSpot assay.

### Memory T cell FluoroSpot assay

To assess memory T cell responses to SARS-CoV-2 in convalescent COVID-19 patient samples we utilized Human IFN-γ/IL-2/TNF FluoroSpot kit (Mabtech), which allows for the detection of T cells producing IFN-γ, IL-2 and/or TNF following stimulation. Cryopreserved PBMCs were thawed at 37°C in RPMI-1640 medium supplemented with 2 mM L-glutamine, 1% penicillin/streptomycin and 10% FCS (all from Thermo Fisher Scientific) and rested for 24 hours at 37 °C and 5% CO_2_. Cell viability after resting was assessed by trypan blue exclusion, and cells were counted using an automated cell counter Countess II (Thermo Fisher Scientific). Viability of > 60% was maintained after the overnight rest. FluoroSpot plates were blocked for 30 minutes with 10% FCS containing RPMI media prior to the addition of 150,000 – 300,000 PBMCs/well. The following stimulation solutions were then added: (i) SNMO peptide pool, (ii) S1-spanning overlapping peptide library, and (iii) N-spanning overlapping peptide library. All wells with SARS-CoV-2 peptide pool stimulations contained the final concentration of 2 μg/mL of peptides and anti-CD28 antibody diluted at 1:100. Negative control contained 0.8% DMSO and anti-CD28 (at 1:100), while positive control contained anti-CD3 and anti-CD28 (both at 1:100). FluoroSpot plates were then incubated for 21 hours at 37 °C and 5% CO_2_ and developed following the manufacturer’s instructions. Fluorescent spots indicating cells that secreted IFN-γ (FITC), IL-2 (Cy3) and/or TNF (Cy5) were detected with IRIS FluoroSpot reader and counted with Apex software (Mabtech). Numbers of responding cells in negative controls were subtracted from stimulated samples to account for background responses. Positivity threshold was set based on pre-pandemic control samples (i.e., average number of polyfunctional T cells + 3 x standard deviation). Average relative spot volume was measured using Apex software to determine the relative amount of each cytokine secreted by cells responding with 3, 2 or 1 cytokine following stimulation.

### Multiplex

Several soluble analytes including CXCL13, IL-6, BAFF, IL-2, IL-21 and IL-15 were measured in plasma diluted 1:2 from acute COVID-19 patients and acute healthy controls using a custom designed multiplex Magnetic Luminex Screening assay (R&D Systems), according to the manufacturer’s instructions.

### Statistics

Statistical analyses were performed using GraphPad Prism (GraphPad Software). Data sets were analyzed using nonparametric Mann Whitney, Wilcoxon matched-pairs signed rank test, Kruskal-Wallis, Friedman or Spearman rank tests. Dunn’s multiple comparisons test was used to correct for multiple comparisons where applicable. Any p values <0.05 were considered to be statistically significant (*p < 0.05; **p < 0.01; and ***p < 0.001). Correlation and hierarchical clustering analysis were performed with GraphPad Prism (v9.0.1) or R (v4.0.2) and RStudio (v1.3.1056) using packages corrplot (v0.84) and heatmaply (v1.1.1).

## Acknowledgements

We would like to thank all donors, health care personnel and members of the Karolinska KI/K COVID-19 Study Group for their participation in the study. The Karolinska KI/K COVID-19 Study Group are as follows: Akber M, Aleman S, Berglin L, Bergsten H, Björkström NK, Brighenti S, Brownlie D, Buggert M, Butrym M, Chambers BJ, Chen P, Cornillet M, Cuapio A, Diaz Lozano I, Dillner L, Dzidic M, Emgård J, Eriksson LI, Flodström-Tullberg M, Färnert A, Gao Y, Glans H, Gorin JB, Gredmark-Russ S, Grip J, Haroun-Izquierdo A, Henriksson E, Hertwig L, Kalsum S, Kammann T, Klingström J, Kokkinou E, Kvedaraite E, Ljunggren HG, Loreti MG, Lourda M, Maleki KT, Malmberg KJ, Marquardt N, Maucourant C, Mårtensson J, Michaëlsson J, Mjösberg J, Moll K, Muvva JR, Nauclér P, Norrby-Teglund A, Palma Medina LM, Parrot T, Perez-Potti A, Persson BP, Radler L, Ringqvist E, Rivera-Ballesteros O, Rooyackers O, Sandberg JK, Sandberg JT, Sekine T, Sönnerborg A, Sohlberg E, Soini T, Strålin K, Svensson M, Tynell J, Unge C, Varnaite R, von Kries A and Wullimann D.

## Funding

S.G-R. was supported by grants from Marianne and Marcus Wallenberg Foundation, and by grants provided by Region Stockholm (ALF project), by Region Stockholm (clinical research appointment), by Center for Innovative Medicine, Region Stockholm and Karolinska Institutet. J.T.S. and R.V. were supported by the Karolinska Institutet PhD Student Funding. H-G.L. and the Karolinska COVID-19 Study Group were supported by the Knut and Alice Wallenberg Foundation and Nordstjernan AB. J.K. was supported by the Swedish Research Council, and Karolinska Institutet Foundations, Karolinska Institutet.

## Author contributions

Conceptualization, J.T.S., R.V., K.B., and S.G-R; Methodology, J.T.S., R.V., and W.C.; Validation, J. T.S., R.V., and W.C; Formal analysis, J.T.S., R.V., K.B., and S.G-R; Investigation, J.T.S., R.V., W.C., K.T.M., M.G., and K. S; Resources, M.B., E.F., M.S., G.A., M.D., L.F., M.S., L. I.E., O.R., A.S., N.K.B., S.A., K.S., H-G.L., P.C., and J.R.M; Data curation, J.T.S., R.V., K.S., N.B., and P.C.; Writing – original draft, J.T.S. and R.V.; Writing - Review & Editing, J. T.S., R.V., S.G.R., K.B., H-G.L., J.K., and all other authors; Visualization, J.T.S., R.V., K.B. and S.G-R; Supervision, S.G-R., K.B., H-G.L., and J.K; Project administration, J.T.S., R.V., K. B., S.G-R., H-G.L., J.K., P.C., S.A., K.S., and N.K.B.; Funding acquisition, H-G.L., S.G-R., and J.K.

## Notes

### Competing Interest Statement

Marcus Buggert is a consultant for Oxford Immunotech.

