## Supplemental figures for "Longitudinal characterization of humoral and cellular immunity in hospitalized COVID-19 patients reveal immune persistence up to 9 months after infection"

**Table of contents**

**Figure S1** Clinical chemistry values in moderate and severe COVID-19 patients during acute disease and at 5-month convalescence. Related to Figure 1.

**Figure S2** Memory B cell and T cell FluoroSpot data, including IgA-mASC data and representative well images from the assay. Related to Figure 4 and Figure 5.

**Figure S3** Flow cytometry gating strategy for circulating T follicular helper cells and antibody-secreting cells. Related to Figure 2.

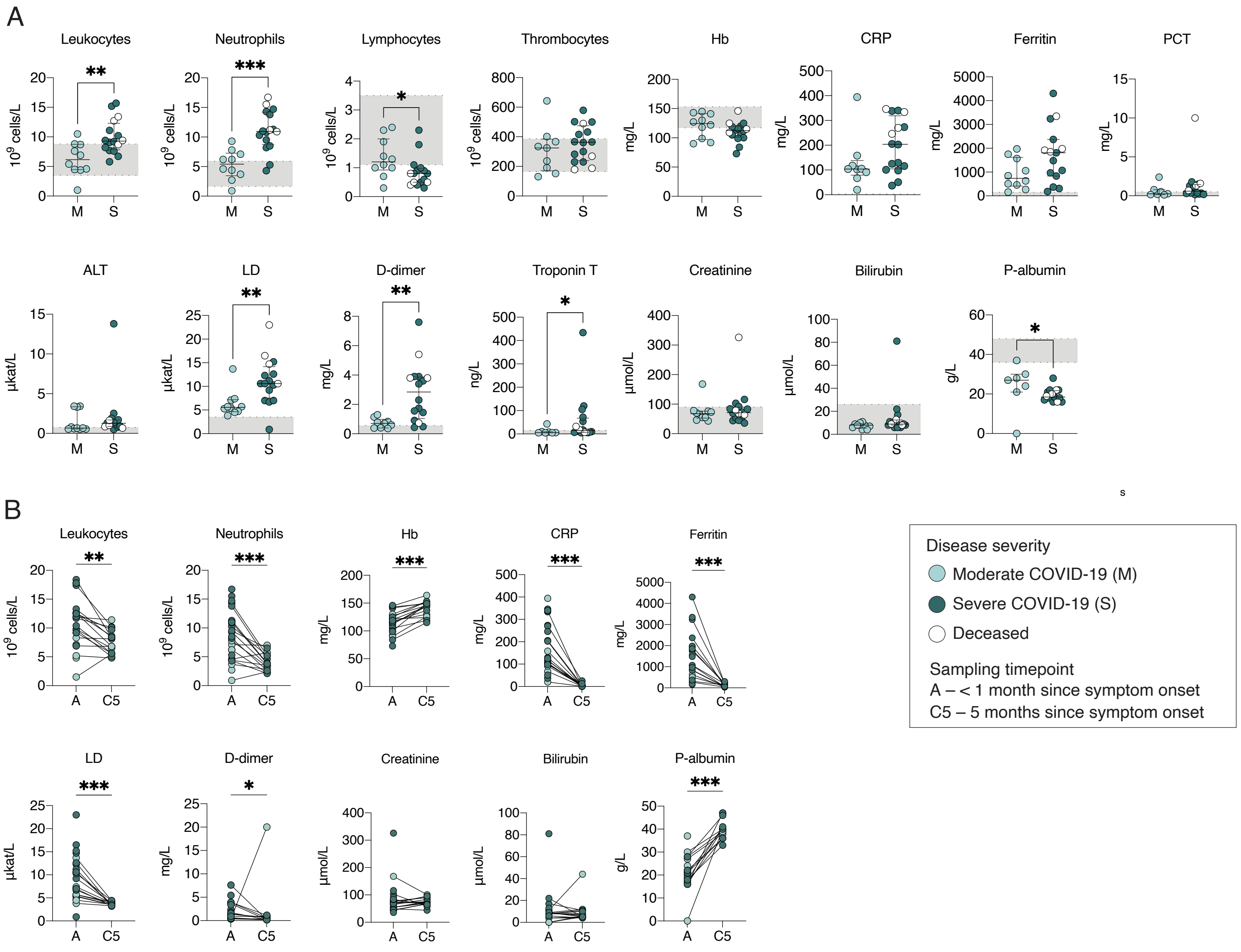


**Supplementary Figure 1. Clinical chemistry values in moderate and severe COVID-19 patients during acute disease and at 5-month convalescence.** (A) Comparison of cell counts, inflammation and organ damage marker levels in peripheral blood between moderate and severe COVID-19 patients during the acute phase. (B) Comparison of clinical chemistry values measured in patients during acute COVID-19 and at 5-month convalescence. Gray boxes indicate the range for reference values. Statistical significance in (A) was assessed by non-parametric Mann-Whitney test. Statistical significance in (B) was assessed by paired, non-parametric Wilcoxon matched-pairs signed rank tests. *p < 0.05, **p < 0.01, and ***p < 0.001.

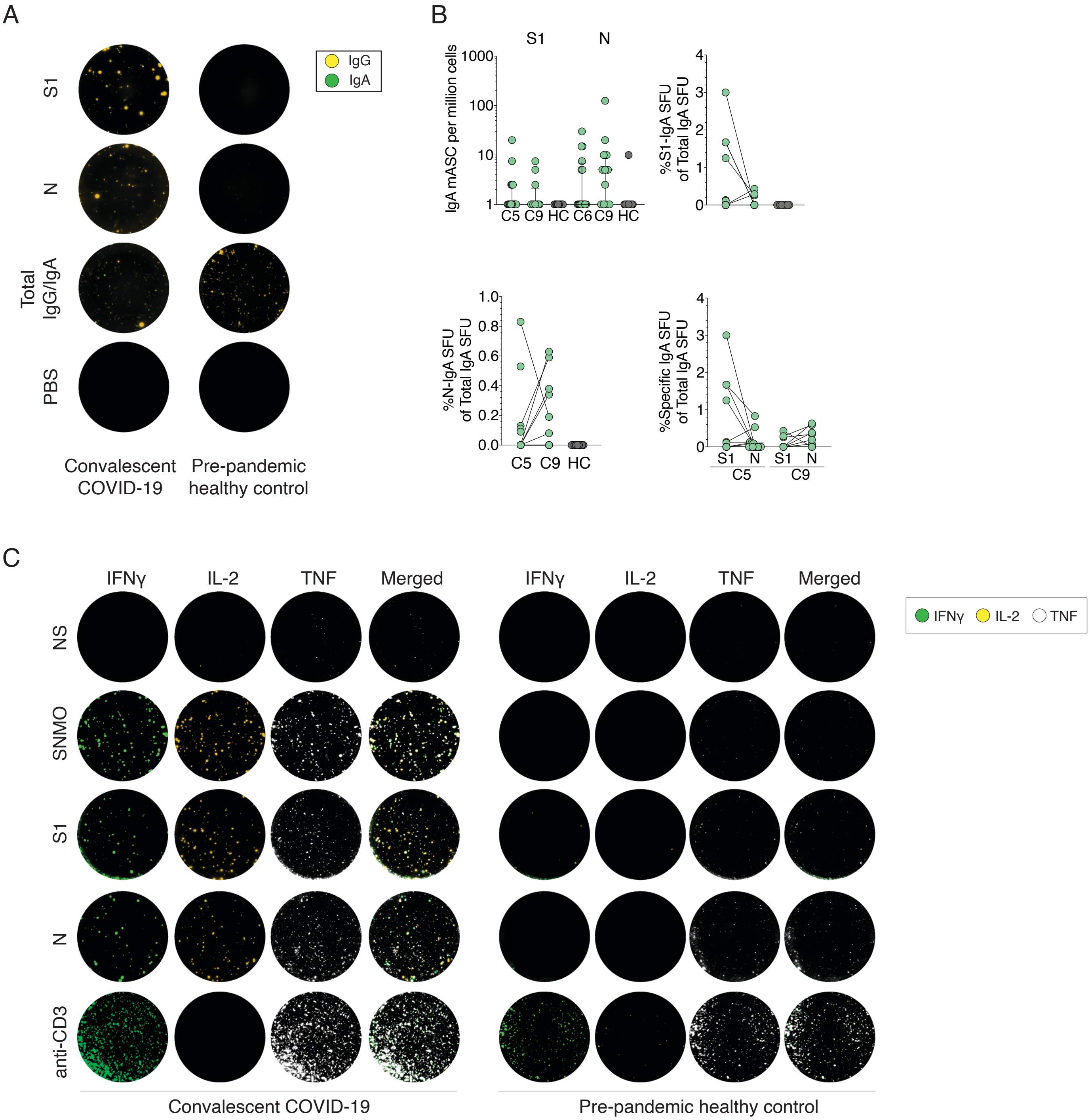


**Supplementary Figure 2. Memory B cell and T cell FluoroSpots.** (A) Representative wells of S1- and N-specific or total IgG and IgA mASCs from a COVID-19 patient at 9-month convalescence and a pre-pandemic healthy control. (B) IgA mASCs per million PBMCs at 5- (C5) and 9-month (C9) convalescence also representated as percent of total IgA mASCs. The final plot compares S1- and N-specific IgA mASCs as percent of total IgA mASCs at C5 and C9. (C) Representative wells from memory T cell FluoroSpot assay from a COVID-19 patient at 9-month convalescence and a pre-pandemic healthy control, either non-stimulated (NS), stimulated with SNMO, S1, N SARS-CoV-2 peptide pools, or with anti-CD3. Green – IFNγ-secreting cells (FITC detection), yellow – IL-2-secreting cells (Cy3 detection), and white – TNF-secreting cells (Cy5 detection). Color for TNF-positive spots was changed from red to white for visualization purposes.

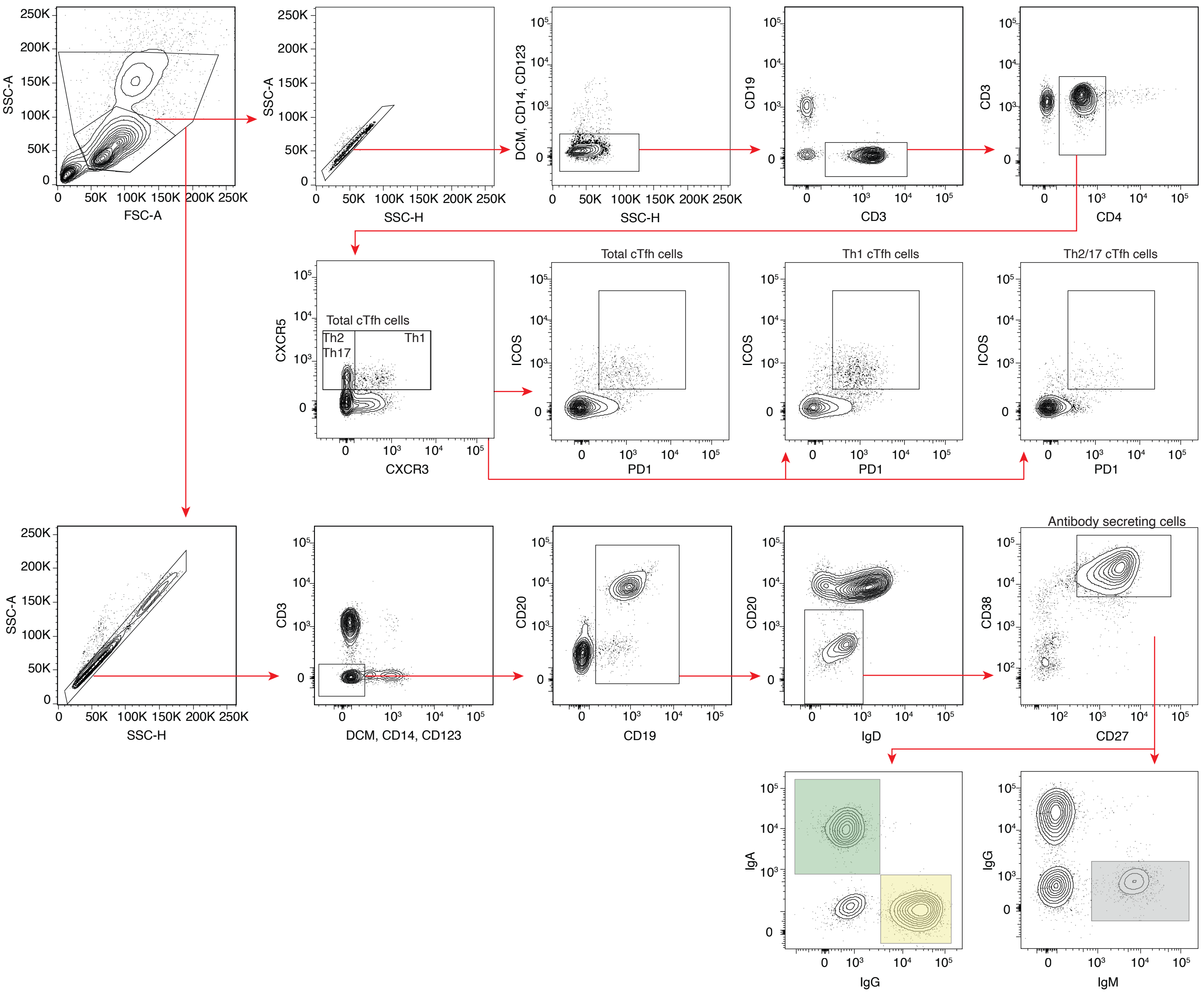


**Supplementary Figure 3. Flow cytometry gating strategy for cTfh cells and antibody-secreting cells.** Activated cTfh cells were defined as ICOS$^{+}$PD1$^{+}$ of (i) total cTfh cells (CD4$^{+}$CXCR5$^{+}$), (ii) Th1-polarized cTfh cells (CXCR5$^{+}$ CXCR3$^{+}$) or (iii) Th2/17- polarized cTfh cells (CXCR5$^{+}$ CXCR3$^{-}$). Antibody-secreting cells were gated from an extended lymphocyte gate and defined as CD38$^{high}$ CD27$^{high}$ cells gated on live CD14$^{-}$ CD123$^{-}$ CD3$^{-}$ CD19$^{+}$ CD20$^{-}$ IgD$^{-}$ cells.
